# Accurate Flow Decomposition via Robust Integer Linear Programming

**DOI:** 10.1101/2023.03.20.533019

**Authors:** Fernando H. C. Dias, Alexandru I. Tomescu

## Abstract

Minimum flow decomposition (MFD) is a common problem across various fields of Computer Science, where a flow is decomposed into a minimum set of weighted paths. However, in Bioinformatics applications, such as RNA transcript or quasi-species assembly, the flow is erroneous, since is obtained from noisy read coverages. Typical generalizations of the MFD problem to handle errors are based on least-squares formulations, or on modeling the erroneous flow values as ranges. All of these are thus focused on error-handling at the level of individual edges.

Interpreting the flow decomposition problem as a robust optimization problem, we lift error-handling from individual edges to *solution paths*. As such, we introduce a new *minimum path-error flow decomposition* problem, for which we give an efficient Integer Linear Programming formulation. Our experimental results reveal that our formulation can account for errors with an accuracy significantly surpassing that of previous error-handling formulations, with computational requirements that remain practical.

## 1 Introduction

### 1.1 Background

In multiassembly problems, multiple genomic sequences in different abundances must be reconstructed from a set of *reads* sequenced from them [30]. Typical examples are the assembly of *RNA transcripts* (or *isoforms* [15, 31], or the assembly of *viral quasi-species* [27, 7]. A standard approach to this problem is first to construct a weighted graph from the reads. For example, in the case of RNA transcripts, one can align the RNA-seq reads to the reference genome and add a graph node for every inferred exon. Each read spanning two exons leads to an edge between the corresponding nodes, directed from smaller to larger genomic positions (obtaining thus a *directed acyclic graph*, or *DAG*). Moreover, edges (or nodes, or both) have an associated weight, e.g. the average read coverage of the corresponding genomic positions. Such a graph is called a *splicing graph*, and it is used by many tools [22]. The multiassembly problem can then be formulated as finding a set of weighted paths whose superposition best “explains” the weights of the graph [30, 24].

If the data is perfect (e.g., no read alignment errors), then the edge weights form a *flow*, in the sense that the sum of the edge weights entering a node equals the sum of the edge weights exiting the same node, for every node different than a source or a sink. A *flow decomposition (FD)* is a set of weighted paths whose superposition matches such flow values, i.e., for every edge, the flow value of the edge equals the sum of the weights of the paths passing through it [1].

Previous research on perfect data noted that flow decompositions of *minimum size* (i.e., with a minimum number of paths, *minimum flow decompositions*, or *MFDs*) largely correspond to correct answers to multiassembly problems [19, 13, 2, 3, 10]. For example, on perfect RNA-seq data, MFDs correspond to the ground truth in over 95% of human genes [19, 13], as we also confirm in this paper. As such, a large amount of theoretical research focused on computing an MFD. While this problem is NP-hard [26] and hard to approximate [11], an exponential-factor approximation exists [16], as well as approximation algorithms for some variants where not all of the flow needs to be decomposed [11], or the flow values can also assume negative values [6]. Also, efficient heuristics [26, 20], or a fixed-parameter tractable algorithm (in the size of the MFD) [13] exist. Recently, it was shown that there exists an Integer Linear Programming (ILP) formulation for MFD [8], which is the fastest practical and exact solution to the problem to date.

### 1.2 Motivation

Since the weights of graphs constructed from real data do not form a flow, practical tools implement generalizations of the MFD problem that account for errors in the edge weights. The most popular one, *least-squares flow decomposition*, asks to find a set of weighted paths that minimise the sum of the squared difference between the weight of each edge and the sum of the weights of the paths passing through it (we call such difference the *superposition error* of an edge). Among all such sets of paths, one with the minimum number of paths is preferred. Many RNA transcript assembly tools use this model, see e.g. [10, 15, 25, 5, 24]. Another more recent variant (*minimum inexact flow decomposition*) is to assume that for each edge, we have an associated interval, and the sum of the weights of the paths passing through the edge must belong to this interval; analogously, among all such solutions, we prefer one of minimum size [28].

These two problems were first solved heuristically. For example, in the case of the former problem, the weights can be first “minimally corrected” so that they become a flow, and then this flow can be heuristically decomposed [25, 10, 5, 15]. For the latter problem, one can compute a “good flow” falling inside the edge intervals and then heuristically decompose this flow. Recently, exact and efficient ILP-based solutions for both problems were shown to exist [8].

Even though practical tools may include additional information to the problem statements (e.g., long reads, as in [31], or pre-assembled contigs as in [2, 18]), models similar to least-squares and inexact flow have been central for dealing with weights not forming perfect flows. In this paper, for the first time, we experimentally evaluate the accuracy of *exact* solutions to the least-squares and minimum inexact flow decomposition models by implementing the exact ILP-based solutions from [8]. On a version of the dataset of [20] where we add errors, we show that the former model reaches an accuracy of 68–80% (depending on how stringent the evaluation metric is and how erroneous the edge weights are), while the latter one reaches an accuracy of 81–91%. On perfect flows, both models are equivalent to MFD, which achieves an accuracy of 95–96%. These results suggest a large gap between our computational models for erroneous and perfect flows.

### 1.3 Contributions

In this paper, we propose a new formulation for the flow decomposition problem for dealing with edge weights not forming a flow. We obtain this by interpreting the problem as a robust optimization problem [21], where it is assumed that some decision variables are prone to errors. A common approach is to model such errors as enlarged bounds for the variables [4]. One can interpret the minimum inexact flow decomposition problem as an instance of this approach. Another approach is to include the errors in the objective function. One can interpret the least-squares flow decomposition problem as an instance of this approach. However, these two previous approaches attach errors to *each edge*. From our point of view, this limits the robustness analysis to a *microscopic* management of errors focused on individual edges.

In this paper, we propose a *macroscopic* management of errors by attaching an error to each *solution path* instead of each edge. In particular, for each solution path, we introduce an error variable and define the *minimum path-error flow decomposition problem* as the problem of finding a set of weighted paths with associated error variables, such that the superposition difference of each edge is within the sum of the error variables of the paths using the edge. Among all solutions to this problem, we prefer minimizing the total path errors and having a minimum number of paths.

Intuitively, as opposed to the previous two formulations, if an edge belongs to multiple solution paths, then each such path contributes with an allowed error since it is more likely that the observed weight of that edge is noisier if it arises by summing up the read coverage of multiple genomic sequences. However, in the inexact flow formulation, the superimposed paths had to belong to a fixed interval, no matter how many paths passed through that edge. Likewise, in the least-squares formulation, the superposition error of an edge contributes the same amount to the objective function no matter how many solutions paths use the edge.

When the graph weights form a flow, this problem is equivalent to MFD; hence it is NP-hard. However, we show that it can be easily and efficiently solved via an ILP formulation, with a quadratic number of variables and constraints in the graph size and the number of solution paths, by adapting the ILP formulations from [8]. In the above-mentioned experimental setting, our formulation achieves an accuracy of 89–95%, just a few percentage points behind the MFD formulation for perfect flows, and running on average in 3 seconds per input weighted graph, and, at the same, each instance was solved in under 7.5 minutes. In the most demanding evaluation scenario, we obtain an accuracy improvement of over 15% over the best previous formulation handling errors. Therefore, we hope that this problem formulation, enhanced with various additional information employed by practical multiassebly tools (such as long reads, as in [31], and also statable into ILP [8]), could lie at the core of more accurate multiassembly tools.

The paper is organized as follows. We review the basic concepts of flow network and flow decomposition and define the problems addressed as well as previous methods attempted for flow decomposition under imperfect flow conditions in Sections 2.1 to 2.3. We next present the robust formulation in Section 3. Numerical experiments are provided in Section 4, and concluding remarks are discussed in Section 5.

### 1.4 Preliminary notions

A directed graph (or simply, a *graph*) is a tuple (*V, E*), with *V* the set of nodes, and *E* ⊆ *V* × *V* the set of edges.^1^ In this paper, we work only with directed acyclic graphs (*DAGs*), that is, graphs containing no directed cycles. We also assume that the DAG has a single *source (s)*, namely a node without incoming edges, and a single *sink (t)*, namely a node without any outgoing edges. A path from *s* to *t* is called an *s-t path*. For a path *P* and an edge (*u, v*) ∈ *E*, we let *P* (*u, v*) = 1, if the edge (*u, v*) belongs to *P*, and *P* (*u, v*) = 0, otherwise.

We also assume to have a *flow value f*_*uv*_ ∈ ℤ^+^ associated to every edge (*u, v*) ∈ *E*, such that the following *flow conservation* condition holds:

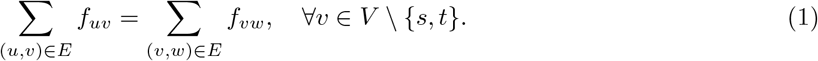

Given such a DAG (*V, E*), with associated flow values *f*, the tuple *G* = (*V, E, f*) is said to be a *flow network*. Given a flow network *G*, a set of *k s*-*t* paths (𝒫 = (*P*_1_, …, *P*_*k*_)) with associated weights *w* = (*w*_1_, …, *w*_*k*_), where each *w*_*i*_ ∈ ℤ^+^, is said to be a *k-flow decomposition* of *G*, if the following *flow superposition* condition holds:

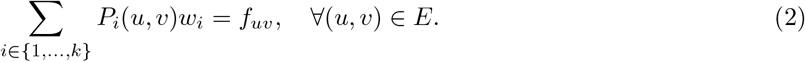

The number *k* of *s*-*t* paths is called the *size* of the flow decomposition 𝒫 and is also denoted |𝒫 |. See Figure 1 for an example of a flow network and of a flow decomposition of it.

**Figure 1:**
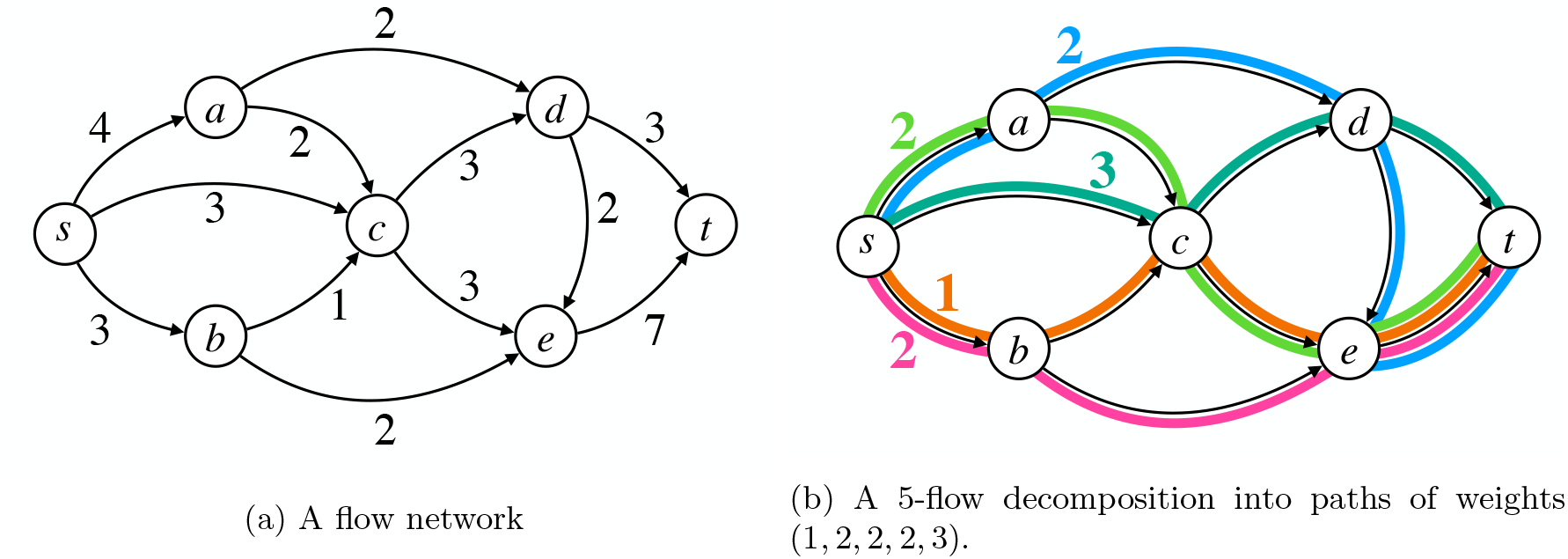
Example of a flow network and a flow decomposition into 5 *s*-*t* paths.

The minimum flow decomposition problem [26] asks to find a flow decomposition of a given flow network of minimum size. Formally, it is defined as follows.

#### Problem 1

(Minimum Flow Decomposition). *Given a flow network G* = (*V, E, f*), *find a minimum-size set of s-t paths 𝒫* = (*P*_1_, …, *P*_*k*_) *with associated weights* (*w*_1_, …, *w*_*k*_), *with each w*_*i*_ ∈ ℤ^+^, *namely, a solution to the following model:*

Minimise |𝒫|

Subject to:

Flow Superposition Condition (2).

Problem 1 admits an efficient Integer Linear Programming formulation, as shown in [8], which works by iterating over *k* in increasing order and checking whether a flow decomposition of size *k* exists.

## 2 Previous approaches for handling errors

In this section, we review three state-of-the-art approaches for decomposing a flow network when the flow values associated with the edges do not satisfy flow conservation anymore (e.g., are arbitrary weights).

### 2.1 Bounded-error Formulation

Let (*V, E*) be a DAG with unique source *s* and unique sink *t*, with an arbitrary value *f*_*uv*_ associated to each edge (*u, v*) ∈ *E* (i.e., not necessarily satisfying the flow conservation condition). The tuple *G* = (*V, E, f*) is called an *imperfect flow network* [8]. Moreover, assume that we are also given an *error bound B*, which intuitively represents the bound by which the Flow Superposition Condition (2) can be relaxed; see Figure 2a for an illustration.

**Figure 2:**
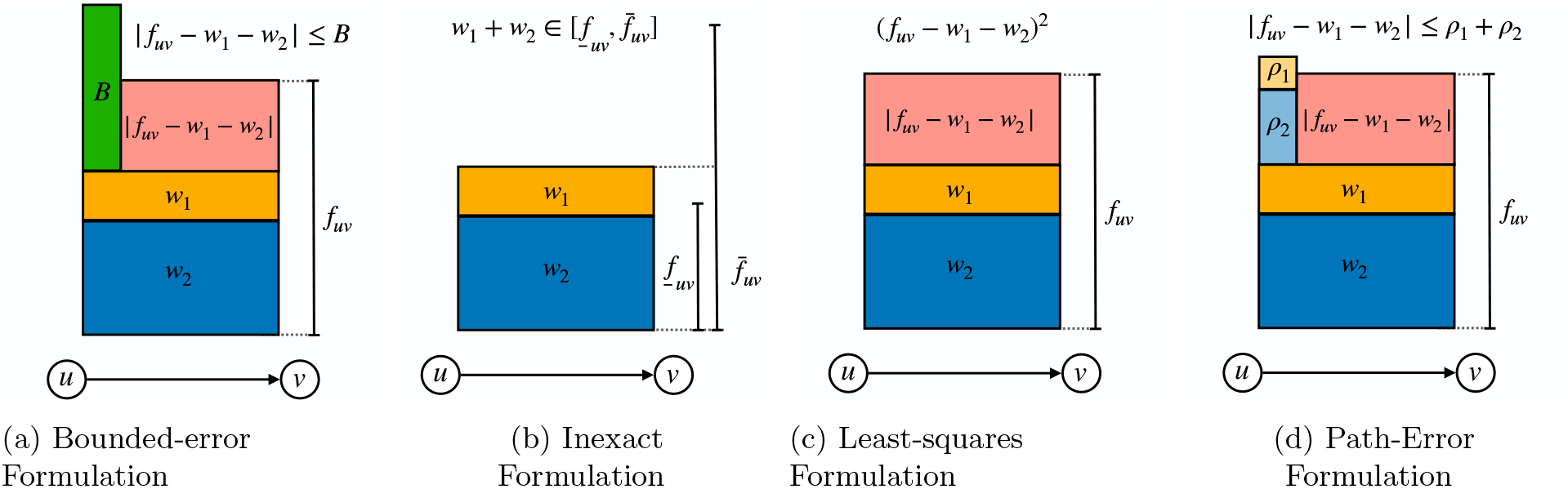
Illustration of the behaviour expected for all methods present in this manuscript. In orange and blue, we show the weights *w*_1_ and *w*_2_ of paths *P*_1_ and *P*_2_, respectively, which we assume to pass through the edge (*u, v*). In red, we show the superposition error, namely, the absolute difference |*f*_*uv*_ − *w*_1_ − *w*_2_|. Each method uses different techniques to control the amount of superposition error. In Figure 2a, an error bound *B* (in green) limits the superposition error; in Figure 2b, superposition error is allowed as long as *w*_1_ + *w*_2_ falls in a given range of the edge (*u, v*); in Figure 2c, a minimization of least-squares of the superposition errors is applied. Finally, in the path-error formulation of Figure 2d, the superposition error is not controlled by a global bound or by the edge (*u, v*) (as in Figures 2a and 2b), but by the sum of paths slacks *ρ*_1_ + *ρ*_2_ of the paths *P*_1_ and *P*_2_, in light orange and light blue, respectively.

Formally, we would like to find a minimum-size set of *s*-*t* paths (*P*_1_, …, *P*_*k*_) with associated weights (*w*_1_, …, *w*_*k*_) such that the following *imperfect flow superposition* condition, with error bound *B*, holds:

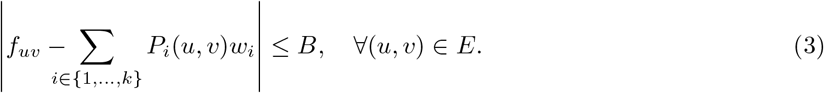

We call the left-hand term of the above inequality *superposition error*. Formally, we have the following problem.

#### Problem 2

(Minimum imperfect flow decomposition (bounded-error)). *Given an imperfect flow network G* = (*V, E, f*), *and an error bound B* ≥ 0, *find a set of s-t paths* 𝒫 = (*P*_1_, …, *P*_*k*_) *with associated weights* (*w*_1_, …, *w*_*k*_), *with each w*_*i*_ ∈ ℤ ^+^, *that are a solution to the following model:*

Minimise | 𝒫 |

Subject to:

Imperfect Flow Superposition Condition with error bound B (3).

Problem 2 is clearly NP-hard, since Problem 1 can be reduced to it by additionally setting *B* = 0. It also admits an efficient Integer Linear Programming formulation [8], which analogously checks for the existence of a set of *k* weighed paths in increasing order on *k*.

### 2.2 Inexact Formulation

The above bounded-error formulation, where the error bound is the same for all edges, has been generalized by [28] to allow for an *error range* specific for each edge. More specifically, we can consider that for each edge (*u, v*) ∈ *E*; we have associated two values 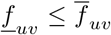. Such a tuple 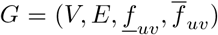 is then called an *inexact flow network* [28].

We now ask for a minimum-size set of *s*-*t* paths (*P*_1_, …, *P*_*k*_) with associated weights (*w*_1_, …, *w*_*k*_) such that the sum of the reported path weights of each edge is in the range 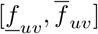; see Figure 2b for an illustration. More precisely, we have the following *inexact flow superposition* condition:

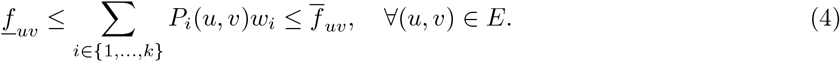

Formally, we have the following problem.

#### Problem 3

(Minimum inexact flow decomposition). *Given an inexact flow network* 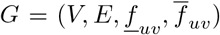, *find a set of s-t paths* 𝒫 = (*P*_1_, …, *P*_*k*_) *with associated weights* (*w*_1_, …, *w*_*k*_), *with each w*_*i*_ ∈ ℤ^+^, *that are a solution to the following model:*

Minimise |𝒫|

Subject to:

Inexact Flow Superposition Condition (4).

Also, Problem 3 is NP-hard, being a generalization of Problems 1 and 2. Likewise, it also admits an efficient Integer Linear Programming formulation [8], which analogously checks for the existence of a set of *k* weighed paths in increasing order on *k*.

### 2.3 Least-squares Formulation

In the previous formulations, the superposition error was bounded by either a global error bound *B* or by a finite range 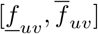. Again, supposing we assume an imperfect flow network. In that case, an alternative is to minimise the sum of all superposition errors. For example, as in a popular least-squares formulation used by [15], minimising:

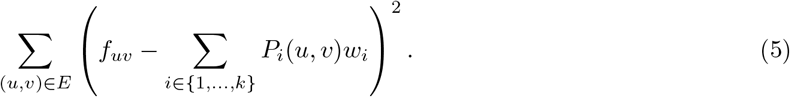

Moreover, overall such sets of *s*-*t* paths (*P*_1_, …, *P*_*k*_), we prefer one of minimum size. Formally, we have the following problem.

#### Problem 4

(Least-squares flow decomposition). *Given an imperfect flow network G* = (*V, E, f*), *find a minimum-size set of s-t* 𝒫 = (*P*_1_, …, *P*_*k*_) *with associated weights* (*w*_1_, …, *w*_*k*_), *with each w*_*i*_ ∈ ℤ^+^, *that are a solution to the following model:*

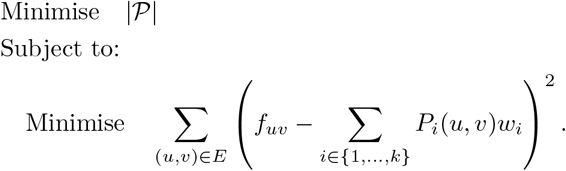

Problem 4 is also NP-hard [24] and can be solved via Integer Quadratic Programming (IQP), see [8] for details. This formulation works by trying all *k* values until |*E*| and taking the set of *k* weighted paths having the least-squared error.

## 3 Minimum Path Error Flow Decomposition

### 3.1 Problem Definition

Assume we have an imperfect flow network and consider the superposition error of an edge (*u, v*). In the bounded and inexact formulations, the superposition error was controlled either by a global error bound or by a range associated to (*u, v*). In both cases, the bound or the range of (*u, v*) stays the same no matter the number of *s*-*t* paths of the decomposition traversing (*u, v*). Moreover, individual edges control the errors in all three previous formulations.

The main idea of our novel path-error formulation is to move the error control mechanism from individual edges to the flow decomposition paths. As such, we can introduce a non-negative *slack* variable *ρ*_*i*_ for each *s*-*t* path *P*_*i*_ and impose that the superposition error of (*u, v*) is at most the sum of the slacks of the *s*-*t* paths using (*u, v*). Intuitively, since the “error” of the path is now associated with paths (and not independently controlled by individual edges), then the fact that a path *P*_*i*_ leads to a substantial superposition error in some edges is overall penalised by increasing the needed slack *ρ*_*i*_ of *P*_*i*_. Formally, we have the following *robust flow superposition* constraint, for each edge (*u, v*) ∈ *E*:

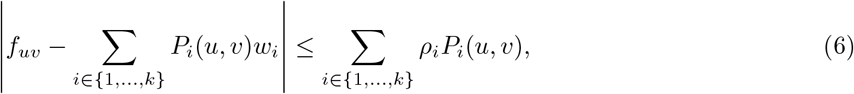

Since the path slacks now control the superposition errors, we can ask for a set of paths of minimum total slack, that is, minimising:

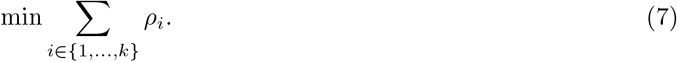

Among all sets of paths of minimum total slack, we prefer one of minimum-size. Formally, we have the following problem.

#### Problem 5

(Minimum Path Error Flow Decomposition). *Given an imperfect network G* = (*V, E, f*), *find a minimum-size set of s-t paths* 𝒫 = (*P*_1_, …, *P*_*k*_) *and associated weights* (*w*_1_, …, *w*_*k*_) *and slacks* (*ρ*_1_, …, *ρ*_*k*_), *with each w*_*i*_ ∈ ℤ^+^, *ρ*_*i*_*x*_*uvi*_ ∈ ℤ^+^ ∪ {0}, *that are a solution to the following model:*

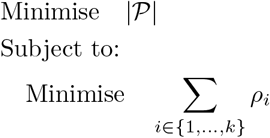

Subject to:

Robust Flow Superposition Condition (6).

To find a solution, we sequentially solve the inner layer in Problem 5 with increasing cardinality of *P*. A minimal path-error flow decomposition is obtained when the accumulated errors from two iterations do not change.

### 3.2 Integer Linear Programming Formulation

In this section, we give an ILP formulation for Problem 5. This follows the same general structure given by [8] for the minimum flow decomposition problem: formulate *k s*-*t* paths, then model the strong flow superposition constraint, and minimise the sum of path slacks. (we call this a *path-error k-flow decomposition*). Moreover, to find a minimum-size path-error flow decomposition, we iterate *k* from 1 to |*E*| and find the smallest such *k* of minimum total slack.

More specifically, for a fixed *k*, to model each path *P*_*i*_, *i* ∈ {1, …, *k*}, we introduce a set of binary variables *x*_*uvi*_ for each edge (*u, v*) ∈ *E*. If *x*_*uvi*_ = 1, the edge (*u, v*) will be part of *P*_*i*_, and if *x*_*uvi*_ = 0, then it is not. Every path *P*_*i*_ is required to start at *s* (i.e., to have exactly one variable *x*_*svi*_ set to 1) and end at *t* (to have exactly one variable *x*_*vti*_ set to 1). Any other node of the graph requires equal in- and out-degree in terms of the binary variables *x*_*uvi*_ (i.e., 0 or 1). More formally, we add the following constraints for every *v* ∈ *V* :

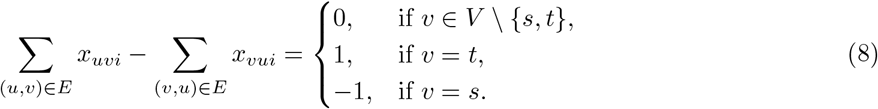

See [8, 23] for a more detailed explanation of why this is a correct formulation of an *s*-*t* path in a DAG.

Alongside these path constraints, we must impose constraint (6). By replacing *P*_*i*_(*u, v*) with *x*_*uvi*_, the following constraint is obtained:

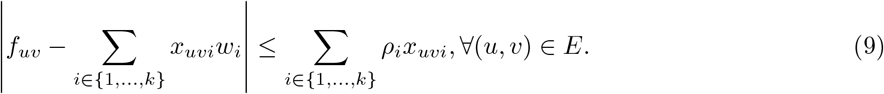

The above constraint contains non-linear terms (*x*_*uvi*_*w*_*i*_ and *ρ*_*i*_*x*_*uvi*_) that standard linear solvers cannot solve. However, using basic linearisation techniques, these terms can be replaced by a set of linear constraints. The product *x*_*uvi*_*w*_*i*_ is a product of two decision variables, one binary and one integer. It can be replaced by a new integer variable *ϕ*_*uvi*_ ∈ ℤ^+^ ∪ {0}, such that:

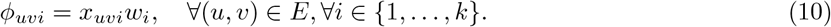

To guarantee equivalence in this substitution of variables, the following additional constraints are necessary. For all (*u, v*) ∈ *E*, and *i* ∈ {1, …, *k*}:

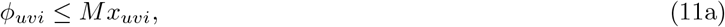

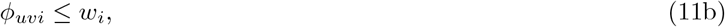

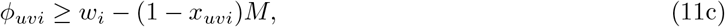

where *M* is a sufficiently large constant so that the inequality in (11a) is always satisfied. Correctness can be seen as follows. If *x*_*uvi*_ = 1, then constraint (11a) holds by the choice of *M*. Constraint (11b) sets *ϕ*_*uvi*_ ≤ *w*_*i*_, and constraint (11c) sets *ϕ*_*uvi*_ ≥ *w*_*i*_; thus *ϕ*_*uvi*_ = *w*_*i*_ = *x*_*uvi*_*w*_*i*_. If *x*_*uvi*_ = 0, then constraint (11a) sets *ϕ*_*uvi*_ ≤ 0. Since *ϕ*_*uvi*_ ≥ 0, we have *ϕ*_*uvi*_ = 0 = *x*_*uvi*_*x*_*uvi*_, and the other two constraints are satisfied.

The terms *ρ*_*i*_*x*_*uvi*_ can be linearised using the same technique by introducing the additional variable *γ*_*uvi*_ ∈ ℤ^+^ ∪ {0} and ensure that *γ*_*uvi*_ = *ρ*_*i*_*x*_*uvi*_. For this case, we will choose (possibly) another significantly large constant *M* that functions as above. For all (*u, v*) ∈ *E*, and *i* ∈ {1, …, *k* }, the resulting constraints are:

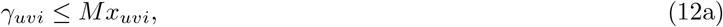

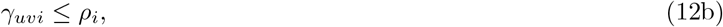

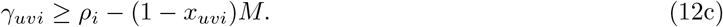

Finally, the absolute value function in the left-hand of the constraint in Equation (6) can also be expanded in the two separated constraints, for all (*u, v*)*inE*:

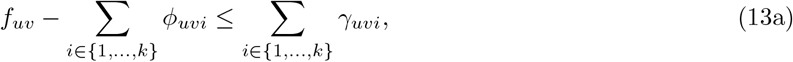

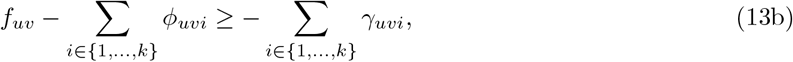

This complete ILP model can be found in Section 6.

## 4 Experiments Design

In this section, we are discussing the solvers implemented in this paper in Section 4.1, the experiments and datasets used to test all formulations in Section 4.2, as well the metrics used to evaluate the formulations and finally present all results in tables in Section 4.3.

### 4.1 Solvers

We implemented all three previous formulations and tested them against our new formulation. We denote the bounded-error formulation (Problem 2) as `Bounded`; the inexact formulation (Problem 3) as `Inexact`; the least-squares formulation (Problem 4) as `LeastSquares`; the minimum path error formulation (Problem 5) as `MinPathError`. We also run the ILP formulation from [8] (Problem 1), which we denote as `MFD`. Our implementations were done using the GUROBI Python API under default settings. We ran our experiments on a personal computer with 16 GB of RAM and an Apple M1 processor at 2.9 GHz. The runtimes of our ILP implementations include the linear scan in increasing order to find the smallest *k* for which there is a solution decomposition (for the formulations that are necessary).

### 4.2 Datasets and metrics

In our experiments, we start from a human transcriptomics dataset generated by [19] containing perfect splice graphs for each human gene. This was built using publicly available RNA transcripts from the Sequence Read Archive with quantification using the tool Salmon [17]^2^. Each input consists of a perfect flow in an acyclic splice graph for each human gene, together with its annotated RNA transcripts (i.e., the ground truth *s*-*t* paths in the graph) and simulated expression values (i.e., ground truth path weights). This dataset was used by e.g. [19, 13] to measure the accuracy of the Minimum Flow Decomposition problem. Additionally, to obtain non-trivial instances, we remove all input graphs that are a single path.

The inputs for `MFD` tests are the perfect inputs from this original dataset. To obtain imperfect flow networks, we introduce errors or noise in the flow values of the edges, as follows. We first assume that the read coverage distribution follows a Poisson distribution, as commonly assumed, see e.g. [15, 5]. For each edge (*u, v*) with perfect flow value 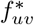, we consider the Poisson distribution Pois 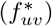. For each allowed error range *ϵ* ∈ {0.25, 0.5, 1}, we compute the (0.5 − *ϵ/*2) ∗ 100-th percentile and (0.5 + *ϵ/*2) ∗ 100-th percentile of this distribution. To get an input for the `Inexact` formulation, we use these two values as the two flow bounds 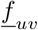 and 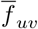, respectively, of edge (*u, v*). As described in [28], it is possible that an infeasible instance is created using this approach for the `Inexact` formulation. In this case, we re-create it until a feasible instance is found. To get an input to be used for the other formulations, for each edge (*u, v*), we set its imperfect flow value *f*_*uv*_ by taking a random sample from the distribution Pois 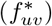, but restricted to the range between these two percentiles (and normalized to the mass of the distribution in this range); note that *ϵ* = 1 corresponds to sampling from the full distribution. The data generation procedure can be found in the https://github.com/algbio/RobustFlowDecomposition, and the new dataset can be found in https://zenodo.org/record/7671871.

As part of our evaluation procedure, we consider three metrics of increasing complexity to indicate if the set of weighted paths output for an input graph by a formulation is a *success*:

**M1:** the superposition of the weighted paths matches the superposition of the ground truth paths (i.e., the original perfect flow);

**M2:** the output paths (as sequences of edges) are exactly the same as the ground truth paths;

**M3:** the output paths (as sequences of edges) are the same as the ground truth paths, and each path has the same weight as the corresponding ground truth path.

For each formulation, we compute *accuracy* as the proportion of all input graphs classified as *success* under each metric. Accuracy computed under the latter two metrics was also used in the evaluation of [19, 13]. We also include the first metric to evaluate how well the formulations can recover the original perfect flow from its erroneous variant (before being split into paths).

### 4.3 Results

In Table 1, we show the accuracy computed over the entire dataset. For baseline values, in this table, we also show the results for `MFD`. Note that, by definition, the value of M1 for `MFD` is 100% since its input flow is perfect, and M1 does not measure how well `MFD` decomposes the flow into paths. Among the formulations handling errors, we observe that `MinPathError` is the best performing under M1, meaning that it can accurately reconstruct the ground truth flow from its erroneous version.

**Table 1:** Proportion of input graphs that are classified as success under each metric. For `MinPathError`, we also list in parentheses the relative improvement of the values of each metric over the corresponding values of best previous formulation handling errors, namely `Inexact`.

| Model | M1 |  |  | M2 |  |  | M3 |
| --- | --- | --- | --- | --- | --- | --- | --- |
| **MFD** | 1.00 |  |  | 0.96 |  |  | 0.95 |

| | $\epsilon = 0.25$ | | | $\epsilon = 0.5$ | | | $\epsilon = 1.00$ | | |
| --- | --- | --- | --- | --- | --- | --- | --- | --- | --- |
| Model | M1 | M2 | M3 | M1 | M2 | M3 | M1 | M2 | M3 |
| **Bounded** | 0.88 | 0.85 | 0.79 | 0.85 | 0.83 | 0.77 | 0.81 | 0.80 | 0.74 |
| **Inexact** | 0.91 | 0.89 | 0.85 | 0.90 | 0.88 | 0.83 | 0.86 | 0.85 | 0.81 |
| **LeastSquares** | 0.80 | 0.78 | 0.74 | 0.80 | 0.76 | 0.75 | 0.75 | 0.72 | 0.68 |
| **MinPathError** | 0.95 | 0.93 | 0.92 | 0.94 | 0.93 | 0.91 | 0.92 | 0.90 | 0.89 |
|  | (+4.30%) | (+4.49%) | (+8.23%) | (+4.45%) | (+5.68%) | (+9.63%) | (+6.97%) | (+5.88%) | (+9.87%) |

Next, we discuss only metrics M2 and M3, which also take into account how well also the ground truth paths (and their weights) are reconstructed. Under these two metrics, `MFD` gives a correct answer in 96– 95% of the inputs, respectively. As errors in the flow values are introduced (i.e. with the increase of *ϵ*), the formulations handling errors show a general decline from these baseline values attained by `MFD`. Among the previous formulations, the best-performing one is `Inexact`. `MinPathError` has a relative improvement over `Inexact` ranging between 4–10%, depending on the metric and the level of errors. For example, at a moderate level of errors (*ϵ* = 0.5), `MinPathError` is only 3–4 percentage points behind `MFD` (recall that `MFD` assumes error-free flows), whereas `Inexact` is 8–10 percentage points behind `MFD`. For the largest level of errors, `MinPathError` presents a 6 percentage points drop in accuracy from the `MFD`, whereas this drop is significantly higher for `Inexact`. Although `Bounded` and `LeastSquares` start with competitive performances with small errors, they drastically drop for moderate and large errors.

In Table 2, we further separate these accuracy results by grouping the inputs based on the number of ground truth paths. As expected, the values of the metrics for all formulations get worse with the increase in the number of ground truth paths. However, compared to `Inexact`, we observe that the results of `MinPathError` are more stable with the increase of this number. For example, on large-error levels (when we sample from the full Poisson distribution), `MinPathError` remains over 82% in all cases, while `Inexact` drops below 70%. This drop is further accentuated for the remaining formulations, i.e., below 60% for both `Bounded` and `LeastSquares`.

**Table 2:**
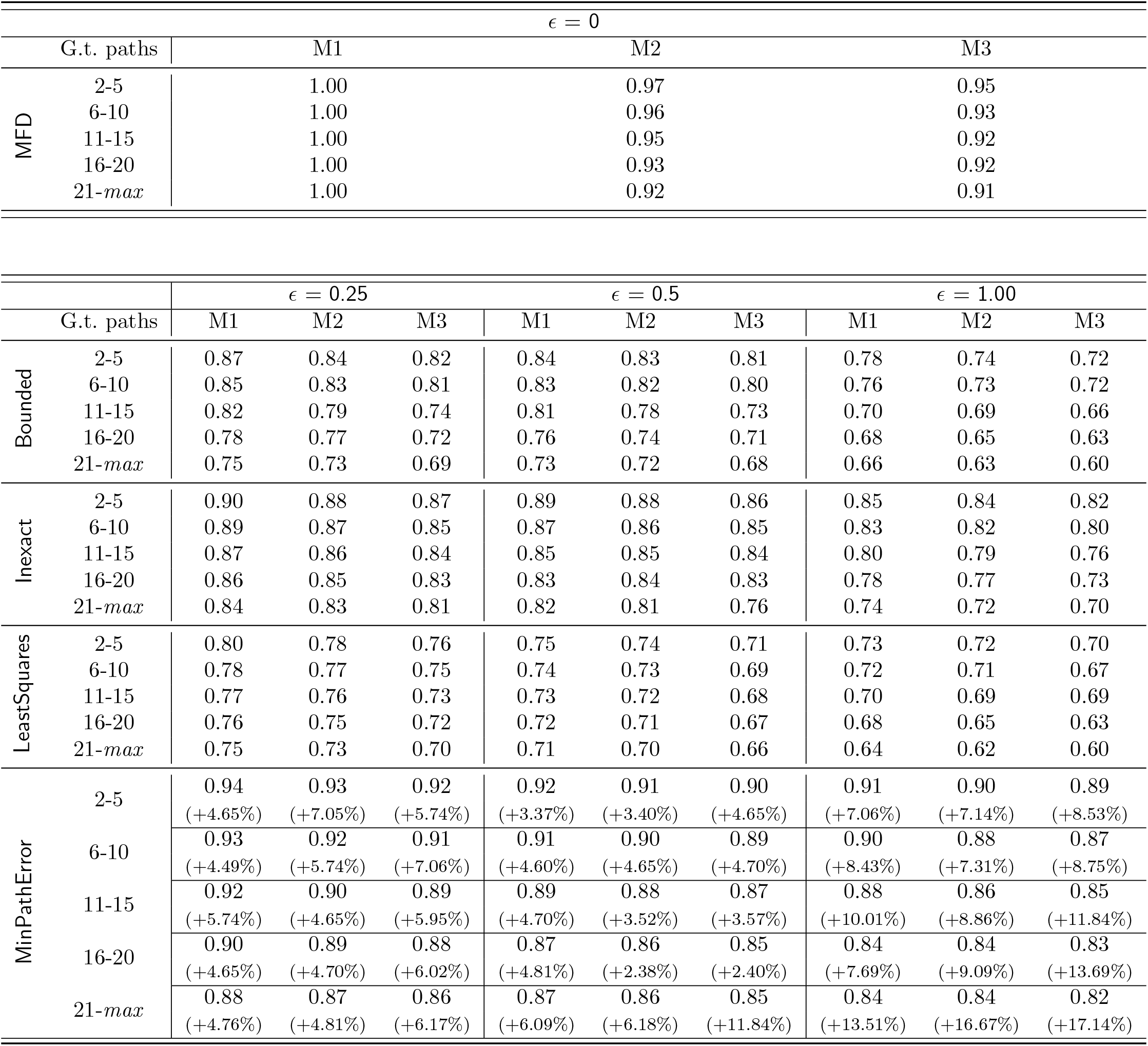
Proportion of input graphs that are classified as success under each metric. Input graphs are partitioned by the number of ground truth paths (column “G.t. paths”). For `MinPathError`, we also list the relative improvement of the values of the metric over the corresponding values of best previous formulation, namely `Inexact`.

In Table 3, we show the runtimes of all four formulations. As expected, `MFD` is the fastest, followed closely by `Inexact`. Thus, `Inexact` is the best among the previous formulations handling errors, being the fastest and most accurate. `MinPathError` is about 4 times slower than `Inexact`, having more variables and including additional auxiliaries due to non-linear components. Nevertheless, across all levels of errors, the average runtime of `MinPathError` is below 5 seconds (the most challenging instances taking at most 450 seconds), and overall it finishes on almost 28 thousand instances within 18 hours, which is practically feasible. The payoff for this larger computational requirement is significantly higher accuracy performance.

**Table 3:**
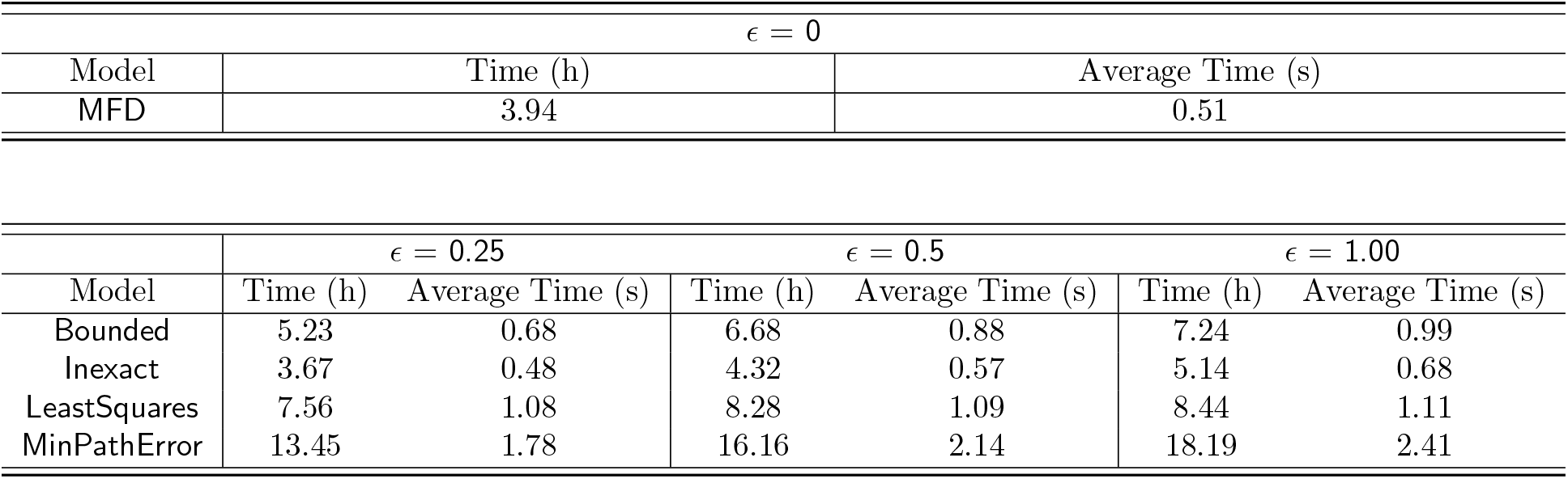
Running times for all formulations tested and implemented in this paper.

It is also important to recall that both `Inexact` and `Bounded` are solved iteratively with increasing *k* until a feasible solution is found. At the same time, `LeastSquares` requires *all* possible values of *k* to be tested, and then an optimal solution is found. Moreover, an optimal solution for `MinPathError` is found when two consecutive iterations do not show improvement in minimizing the combined path error. Because the reported runtime considers all iterations executed for each method, it is crucial to understand why the runtime of the two latter formulations is considerably larger than the former ones.

## 5 Conclusion

By interpreting the imperfect flow decomposition problem as a robust optimization problem, we shifted the management of errors from the microscopic focus on the edges of the graph to the macroscopic focus on the weights of the solution paths. This led to a significant improvement in accuracy concerning the previous error-handling formulations, even on a large amount of errors, being only closely behind decomposing the flow on error-free inputs. This increase in accuracy comes at the cost of an increased running time; however, despite the NP-hardness of the problem, this is still feasible in practice, with any instance finishing in at most 7.5 minutes, while the average runtime over the entire dataset is much smaller.

In this paper, we focused only on giving a flow decomposition formulation better handling errors. In particular, we considered the results of `MFD` on perfect flows to be a gold standard. However, in practical Bioinformatics tools, additional constraints can be added to restrict the solution space further. For example, it is known that if one is also given a set of paths that must appear as subpath in at least one solution path (*subpath constraints*, arising from long reads aligned to the graph), then the accuracy further increases [29]. Such subpath constraints have been added on top of the ILP formulation for `MFD`, see [8]. They can also be trivially added to our ILP from Section 3 by analogously requiring that they appear as subpaths also in our solution paths. We leave this problem for future work since, as already mentioned, this paper focuses on better handling erroneous flows. Overall, given the performance and flexibility of such ILP formulations, they can be at the core of future RNA assemblers.

Although our formulation requires a larger number of variables and constraints to express uncertainty, its size is still asymptotically quadratic, similar to the previous ILP formulations for MFD. Considering the further development of ILP solvers, this will be only a minor limitation in downstream applications. Currently, the main drawback is the dependency of *k*. An improvement of the overall formulation is to obtain a direct estimation of the number *k* of paths of each decomposition which would allow for avoiding multiple iterations. Finally, we look forward to extending different ILP formulations for this problem involving the stochastic formulation [14], uncertainty in cyclic graphs [9], and safety [12].

## Acknowledgment

This work was partially funded by the European Research Council (ERC) under the European Union’s Horizon 2020 research and innovation programme (grant agreement No. 851093, SAFEBIO), partially by the Academy of Finland (grants No. 322595, 352821, 346968).

## 6 Full ILP formulation for Path-Error *k*-Flow Decomposition

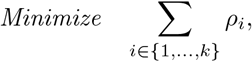

*Subject to:*

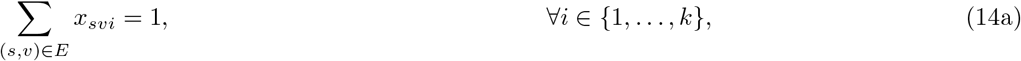

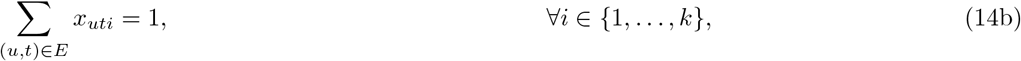

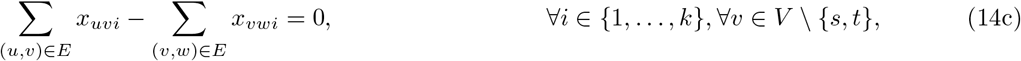

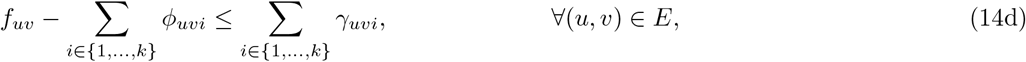

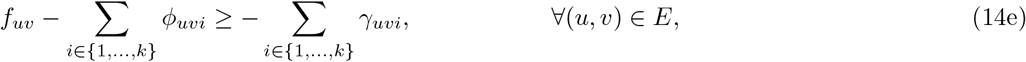

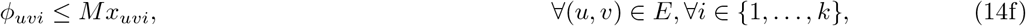

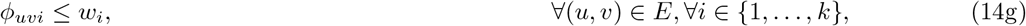

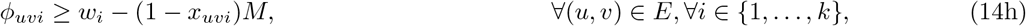

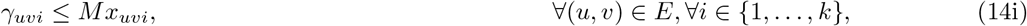

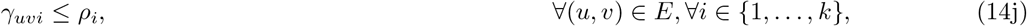

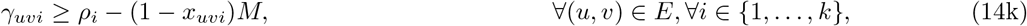

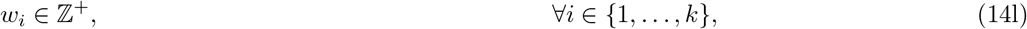

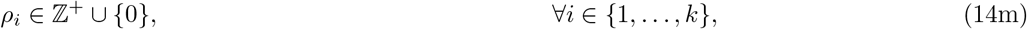

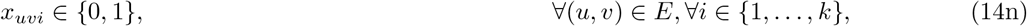

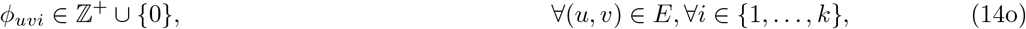

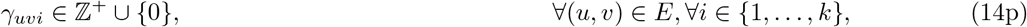

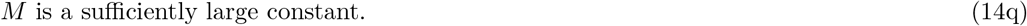

*M* is a sufficiently large constant. (14q)

## Footnotes

1 Since *E* ⊆ *V* × *V*, there can be no *parallel* edges with the same endpoints. However, there is a trivial observation that there is no loss of generality because, for flow decomposition problems, such edges can be subdivided by introducing a new node in the middle of each parallel edge.

2 The full dataset from [20] is available at https://zenodo.org/record/1460998. We use the file rnaseq/sparse_quant_ SRR020730.graph

## Notes

### Competing Interest Statement

The authors have declared no competing interest.

## References

[1] Ravindra K Ahuja, Thomas L Magnanti, and James B Orlin. Network flows. 1988.

[2] Jasmijn A Baaijens, Leen Stougie, and Alexander Schönhuth. Strain-aware assembly of genomes from mixed samples using flow variation graphs. In International Conference on Research in Computational Molecular Biology, pages 221–222. Springer, 2020.

[3] Jasmijn A Baaijens, Bastiaan Van der Roest, Johannes Köster, Leen Stougie, and Alexander Schönhuth. Full-length de novo viral quasispecies assembly through variation graph construction. Bioinformatics, 35(24):5086–5094, 2019.

[4] Aharon Ben-Tal and Arkadi Nemirovski. Robust solutions of linear programming problems contaminated with uncertain data. Mathematical programming, 88:411–424, 2000.

[5] Elsa Bernard, Laurent Jacob, Julien Mairal, and Jean-Philippe Vert. Efficient rna isoform identification and quantification from rna-seq data with network flows. Bioinformatics, 30(17):2447–2455, 2014.

[6] Manuel Cáceres, Massimo Cairo, Andreas Grigorjew, Shahbaz Khan, Brendan Mumey, Romeo Rizzi, Alexandru I. Tomescu, and Lucia Williams. Width helps and hinders splitting flows. In Shiri Chechik, Gonzalo Navarro, Eva Rotenberg, and Grzegorz Herman, editors, 30th Annual European Symposium on Algorithms, ESA 2022, September 5-9, 2022, Berlin/Potsdam, Germany, volume 244 of LIPIcs, pages 31:1–31:14. Schloss Dagstuhl - Leibniz-Zentrum für Informatik, 2022.

[7] Jiao Chen, Yingchao Zhao, and Yanni Sun. De novo haplotype reconstruction in viral quasispecies using paired-end read guided path finding. Bioinformatics, 34(17):2927–2935, 2018.

[8] Fernando H. C. Dias, Lucia Williams, Brendan Mumey, and Alexandru I. Tomescu. Fast, Flexible, and Exact Minimum Flow Decompositions via ILP. In RECOMB 2022 - 26th Annual International Conference on Research in Computational Molecular Biology, volume 13278 of Lecture Notes in Computer Science, pages 230–245. Springer, 2022.

[9] Fernando HC Dias, Lucia Williams, Brendan Mumey, and Alexandru I Tomescu. Minimum flow decomposition in graphs with cycles using integer linear programming. arXiv preprint arXiv:2209.00042, 2022.

[10] Thomas Gatter and Peter F Stadler. Ryūtō: network-flow based transcriptome reconstruction. BMC bioinformatics, 20(1):1–14, 2019.

[11] Tzvika Hartman, Avinatan Hassidim, Haim Kaplan, Danny Raz, and Michal Segalov. How to split a flow? In 2012 Proceedings IEEE INFOCOM, pages 828–836. IEEE, 2012.

[12] Shahbaz Khan, Milla Kortelainen, Manuel Cáceres, Lucia Williams, and Alexandru I Tomescu. Safety and completeness in flow decompositions for RNA assembly. In International Conference on Research in Computational Molecular Biology, pages 177–192. Springer, 2022.

[13] Kyle Kloster, Philipp Kuinke, Michael P O’Brien, Felix Reidl, Fernando Sánchez Villaamil, Blair D Sullivan, and Andrew van der Poel. A practical fpt algorithm for flow decomposition and transcript assembly. In 2018 Proceedings of the Twentieth Workshop on Algorithm Engineering and Experiments (ALENEX), pages 75–86. SIAM, 2018.

[14] Can Li and Ignacio E Grossmann. A review of stochastic programming methods for optimization of process systems under uncertainty. Frontiers in Chemical Engineering, 2:34, 2021.

[15] Wei Li, Jianxing Feng, and Tao Jiang. IsoLasso: a LASSO regression approach to RNA-Seq based transcriptome assembly. Journal of Computational Biology, 18(11):1693–1707, 2011.

[16] Brendan Mumey, Samareh Shahmohammadi, Kathryn McManus, and Sean Yaw. Parity balancing path flow decomposition and routing. In 2015 IEEE Globecom Workshops (GC Wkshps), pages 1–6. IEEE, 2015.

[17] Rob Patro, Geet Duggal, and Carl Kingsford. Salmon: accurate, versatile and ultrafast quantification from RNA-seq data using lightweight-alignment. BioRxiv, page 021592, 2015.

[18] Mihaela Pertea, Geo M Pertea, Corina M Antonescu, Tsung-Cheng Chang, Joshua T Mendell, and Steven L Salzberg. StringTie enables improved reconstruction of a transcriptome from RNA-seq reads. Nature biotechnology, 33(3):290–295, 2015.

[19] Mingfu Shao and Carl Kingsford. Accurate assembly of transcripts through phase-preserving graph decomposition. Nature biotechnology, 35(12):1167–1169, 2017.

[20] Mingfu Shao and Carl Kingsford. Theory and a heuristic for the minimum path flow decomposition problem. IEEE/ACM transactions on computational biology and bioinformatics, 16(2):658–670, 2017.

[21] Allen L Soyster. Convex programming with set-inclusive constraints and applications to inexact linear programming. Operations research, 21(5):1154–1157, 1973.

[22] Stefan Stamm, Shani Ben-Ari, Ilona Rafalska, Yesheng Tang, Zhaiyi Zhang, Debra Toiber, TA Thanaraj, and Hermona Soreq. Function of alternative splicing. Gene, 344:1–20, 2005.

[23] Leonardo Taccari. Integer programming formulations for the elementary shortest path problem. European Journal of Operational Research, 252(1):122–130, 2016.

[24] Alexandru I Tomescu, Travis Gagie, Alexandru Popa, Romeo Rizzi, Anna Kuosmanen, and Veli Mäki-nen. Explaining a weighted dag with few paths for solving genome-guided multi-assembly. IEEE/ACM transactions on computational biology and bioinformatics, 12(6):1345–1354, 2015.

[25] Alexandru I Tomescu, Anna Kuosmanen, Romeo Rizzi, and Veli Mäkinen. A novel min-cost flow method for estimating transcript expression with RNA-Seq. In BMC bioinformatics, volume 14, pages S15:1–S15:10. Springer, 2013.

[26] B. Vatinlen, F. Chauvet, P. Chrétienne, and P. Mahey. Simple bounds and greedy algorithms for decomposing a flow into a minimal set of paths. European Journal of Operational Research, 185(3):1390–1401, 2008.

[27] Kelly Westbrooks, Irina Astrovskaya, David Campo, Yury Khudyakov, Piotr Berman, and Alex Zelikovsky. HCV quasispecies assembly using network flows. In International Symposium on Bioinformatics Research and Applications, pages 159–170. Springer, 2008.

[28] Lucia Williams, Gillian Reynolds, and Brendan Mumey. RNA Transcript Assembly Using Inexact Flows. In 2019 IEEE International Conference on Bioinformatics and Biomedicine (BIBM), pages 1907–1914. IEEE, 2019.

[29] Lucia Williams, Alexandru Tomescu, Brendan Marshall Mumey, et al. Flow decomposition with subpath constraints. In 21st International Workshop on Algorithms in Bioinformatics (WABI 2021). Schloss Dagstuhl-Leibniz-Zentrum für Informatik, 2021.

[30] Yi Xing, Alissa Resch, and Christopher Lee. The multiassembly problem: reconstructing multiple transcript isoforms from est fragment mixtures. Genome research, 14(3):426–441, 2004.

[31] Qimin Zhang, Qian Shi, and Mingfu Shao. Scallop2 enables accurate assembly of multiple-end rna-seq data. bioRxiv, 2021.

